# Variant Mapping Application: Customize And Annotate Figures Of Voltage-Gated Sodium Channel

**DOI:** 10.1101/2023.12.09.570948

**Authors:** Winnie Wen, Arjun Mahadevan, R. Ryley Parrish, Richard Dean, Alison Cutts, JP Johnson

## Abstract

Voltage-gated sodium ion channels allow for the initiation and transmission of action potentials. There is a high interest in research and drug development to selectively target these ion channels to treat epilepsy and other disorders such as pain. Scientific literature and presentations often incorporate maps of these integral membrane proteins with markers indicating gene mutations to highlight genotype/phenotype correlations. There is a need for automated tools to create high quality figures with mutation (variant) locations displayed on these channel maps. This manuscript introduces a simple application to create visualization for mutations on alpha voltage-gated sodium channels, created using the D3.js library. The application allows for mapping of variant sequences, as well as important properties like the type of variant and the phenotypes linked to the variant. It also allows for customizability and the production of high-quality images for publication. This application and code base can further be extrapolated to other ion channels as well.

## Introduction

### Role of Voltage-Gated Sodium Ion Channels in Physiology

Voltage-gated sodium ion channels are integral membrane proteins that induce the rising phase of the action potential in most electrically excitable cells, including neurons (Catterall, 2000).

The channel first opens, allowing sodium ions to flow into the cell and depolarizing the cell membrane potential. Opening is followed by a conformational change to inactivated states, preventing further sodium influx and allowing repolarization (Armstrong, 2006; Oliva et al., 2012). As a result, gain-or loss-of-function mutations in these ion channels result in abnormal excitable cell activity and leads to varied disorders in the nervous system, skeletal muscle, and cardiac tissue (Fouda et al., 2022; Meisler et al., 2021; Stafstrom, 2007).

### Structure of Sodium Channels

Graphical representations of sodium channels are often incorporated into scientific literature, where they are depicted as having four domains (DI – DIV) with six transmembrane segments (S1-S6) in each (Catterall & Swanson, 2015). Segments S1-S4 compose the voltage sensor region with S4 being the positively charged segment that dynamically shifts in response to depolarization inducing the opening (or inactivation) of the channel and the initiation of (or cessation of) sodium influx (Peters et al., 2016). Segments S5 and S6, with the intervening re-entrant P loop, create the pore forming region (Peters et al., 2016; Sands et al., 2005).

### Related Work

Despite the common occurrence of figures for sodium channels in publications, there are few applications that generate diagrams of these proteins automatically. Many of these diagrams are annotated by hand (Goldberg et al., 2007) or manually created through an image editing or presentation program such as PowerPoint (Catterall, 2012; Meisler & Kearney, 2005; Oliva et al., 2012). The few applications that exist specifically for producing figures of voltage-gated channels require the user to place mutation sequences in position manually or require the data to be in a specific format that is inaccessible and unintuitive to most users. TopDraw (Bond, 2003), for example, allows for easy sketching of protein topologies, but it does not provide the ability to annotate and is solely used to derive a sketch from another source such as HERA. TOPO2 (Johns, 2010) is unable to predict the location of the transmembrane segments and only exports the figure as a PNG, which is a raster image format that is sometimes not accepted by publishers. Similarly, NaView (Afonso et al., 2022) generates a visualization automatically and provides customizability by allowing the user to alter colors and text position, but annotations need to be manually added. Furthermore, user-provided input must be a UniProt formatted string and/or a JSON object, which may be inaccessible or tedious for most users.

In comparison, this Variant Mapping application is a user-friendly D3.js and React.js application that can be used to automatically generate a scaled SVG figure of a sodium channel with annotations for mutation sequences, custom labels, and legends. The application also allows custom color selection, filters, toggles, and sliders that can be used to customize the image before exporting to a PNG or SVG file at resolutions acceptable for journal submission.

## Methods and Results

*Link to Application:* Variant Mapping Application(ionchannel-biology.github.io)

### Technologies Used

The Variant Mapping application primarily uses a combination of D3.js and React.js, which are both JavaScript libraries. D3.js, also known as D3 or Data-Driven Documents, uses web standards like HTML, CSS, and SVG to produce dynamic, interactive, and customized data visualizations. The library is particularly powerful in data manipulation and is extremely fast due to its minimal overhead in that it only modifies the attributes that change. This makes it an ideal choice for mathematical functions, analytical tasks, animations, data manipulation, and scaling of data within applications to create beautiful and powerful visualizations. React.js, also known as React, is the other library used and is often utilized to create interactive websites and build user interfaces through the production of components. Since the D3 community has not yet established a standard way to create components, React is ideal to control the rendering and re-rendering of elements.

While integrating D3 and React can be tricky because they both want to manipulate the DOM (Battle et al., 2021), doing so makes the code declarative instead of imperative (Wattenberger, 2022). The integration also allows for most components to be separated, making the elements reusable and the application is more scalable for future updates.

### Structure and Features

Data on the structure of sodium channels is sourced from UniProt and converted to a ^*^.csv file.

The file contains information on the size of the membrane and paths, as well as the relative locations for markers when mutation sequences are later input. The relationship between the elements in the D3 layout is the same as the relationship between the domains, segments, and loops depicted in current literature. The domains are each a group element, the transmembrane segments are rectangles within each domain group, and the loops are path elements in their respective domains. The loops are also scaled proportional to the extracellular and cytoplasmic regions of the protein. D3 provides an advantage here because the path tag allows for the ability to create scalable curves, whereas plain HTML does not have equivalent functionality (Sweeny, 2018).

There are two main inputs for the user to supply their own data: a form for inputting mutational sequences one at a time (Figure 1) and an input to upload an Excel or CSV file. The option to upload Excel files is more accessible compared to earlier applications, where users only had the option to import UniProt code or JSON objects, since most users will have access to Microsoft Excel. This option also provides a quicker alternative to users to upload data if they wish.

**Figure 1:**
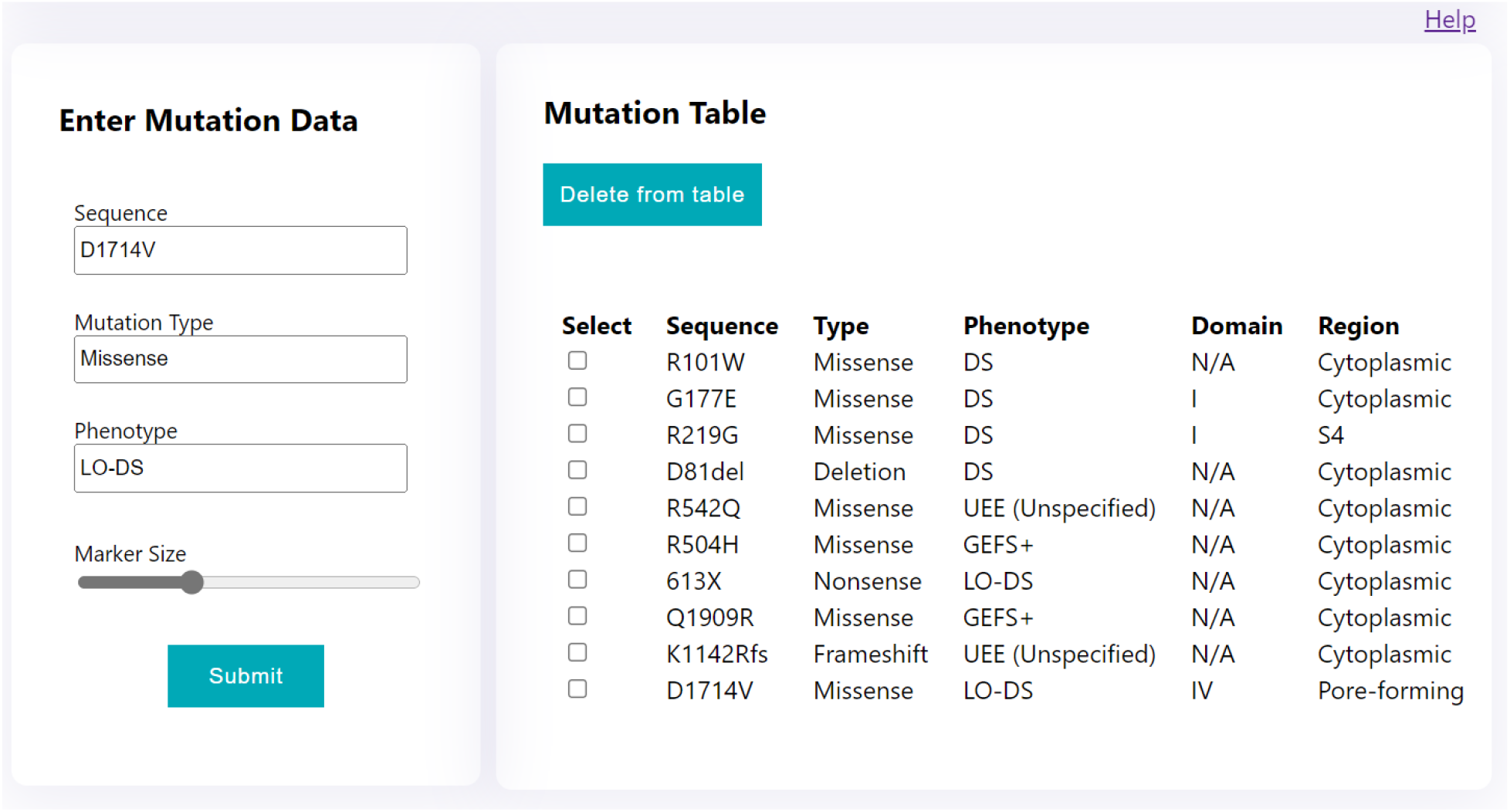
Form and Resulting Table. Users enter mutation data in the form on the left. The table on the right shows an example of what is displayed after entering multiple mutation sequences.

A [colour picker](https://embed.plnkr.co/xP6nHx/), is added for additional customizability and for users to have the flexibility to decide the story they want to tell with their data. For example, users can select darker colours for more severe diagnoses and lighter colours for less severe diagnoses. In another instance, users may select warm colours for loss of function mutations and cool colours for gain of function mutations.

Additionally, the legend has added filters where the user can select the text for the specific mutations they want. This is for added customizability and to provide users with an easier solution to spot patterns between similar types of mutations or diagnoses. Moveable text labels, toggles, and a slider for the legend position have also been added for customizability.

The Variant Mapping application will be hosted on a public server and can be accessed locally on any computer Variant Mapping Application (ionchannel-biology.github.io). The application produces a figure of the desired sodium channel and a table containing the user-input data along with the addition of a domain label and region label (Figure 1). Users can also choose to download the figure as a PNG or SVG (Figure 2) and export the table as an Excel spreadsheet (Figure 3).

**Figure 2:**
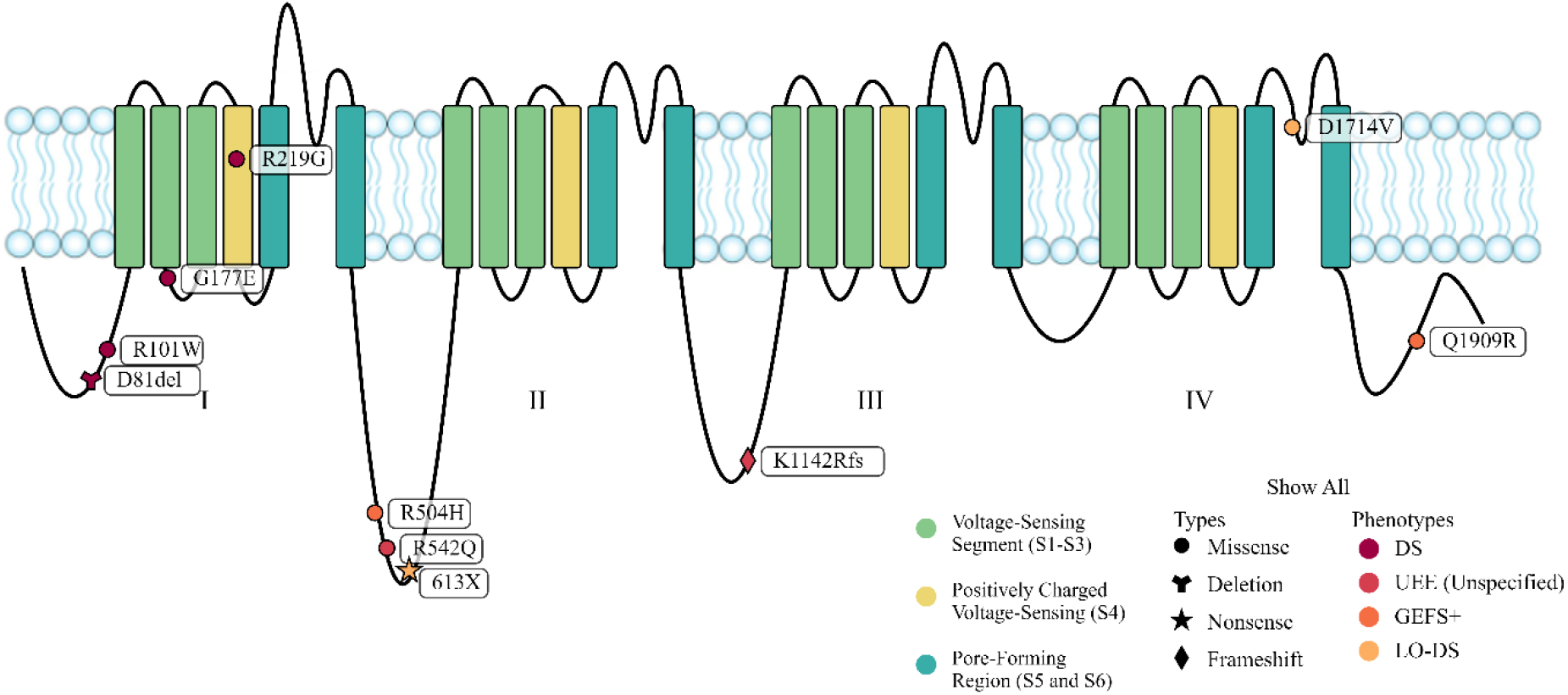
Resulting Figure with Multiple Markers. The figure that is produced when the user exports the diagram as a PNG or SVG. Before exporting, users can customize the figure by moving the labels, altering the colors, filtering the visible mutations, and toggling the legend on/off.

**Figure 3:**
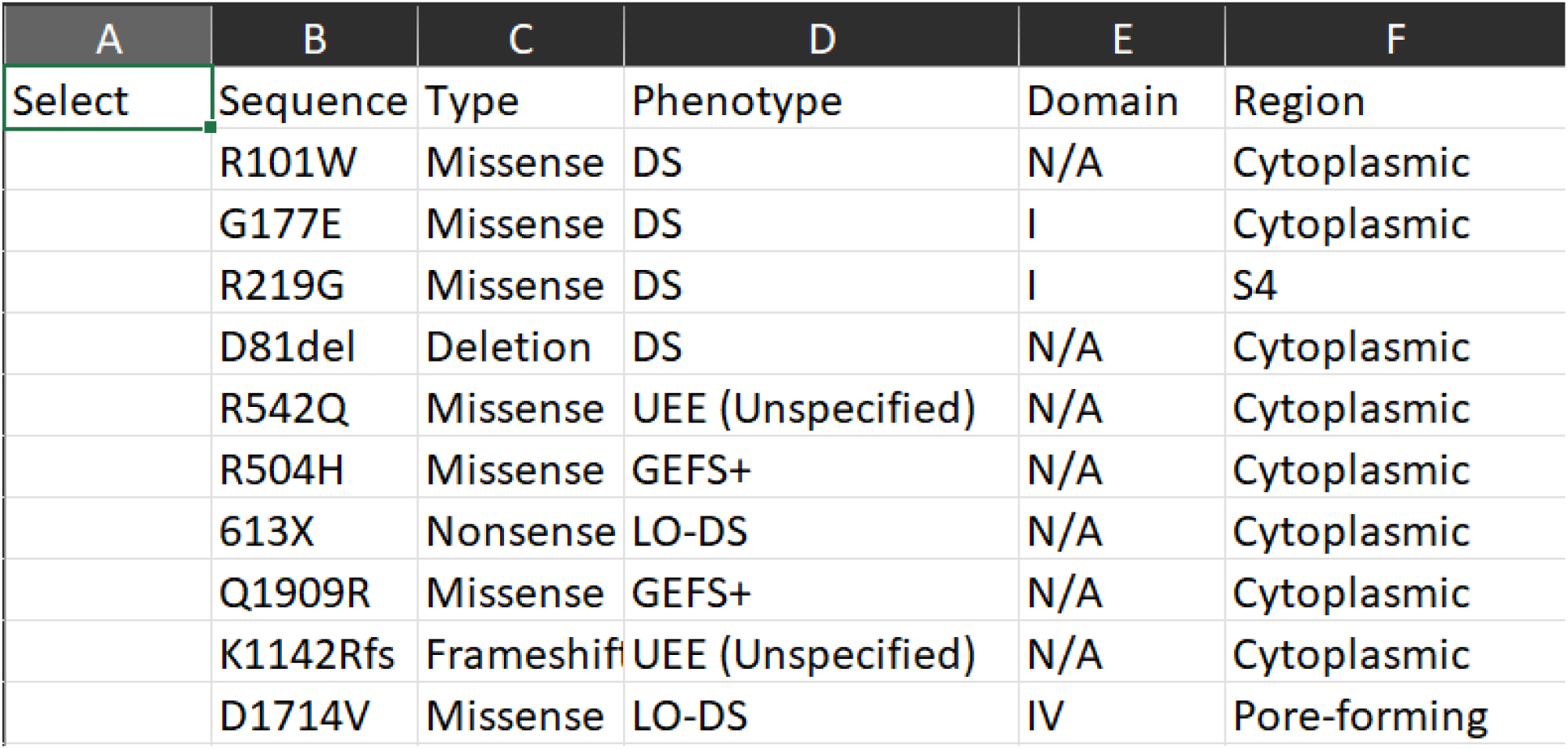
Resulting Excel Spreadsheet. The spreadsheet of the user-inputted mutations after exporting the displayed table to an XLSX file.

## Discussion

In this current work, we demonstrate the Variant Mapping application and are providing public access to the application. We believe this work sets a foundation for generation of other similar tools that not only ease production of high-quality figures but can also be used as a learning tool for understanding ion channel mutations. Future updates could focus on extrapolating to other voltage-gated ion channel families such as calcium and potassium channels. The current code could also be altered to become more flexible by adding the beta subunits of the sodium channels. In summary, we are excited to provide this tool as an open access application to the scientific community and we hope this encourages other similar work that will aid in the advancements in precision medicine and provide a valuable learning tool for students of ion channel function and dysfunction.

## Data availability statement

The raw data supporting the conclusions of this article will be made available by the authors, without undue reservation.

## Acknowledgments

We would like to thank the entire Xenon family for their support of this work.

## Author contributions statement

JPJ conceived this work. WW and AM wrote the code and designed the visualizations. RD and AC provided the raw data for the application and RD, AC, and RRP advised on the project. WW, AM, and RRP wrote the manuscript. All authors edited and approved the final draft.

